# Happy you, happy me: expressive changes on a stranger’s voice recruit faster implicit processes than self-produced expressions

**DOI:** 10.1101/518324

**Authors:** Laura Rachman, Stéphanie Dubal, Jean-Julien Aucouturier

**Affiliations:** Inserm U1127, CNRS UMR 7225, Sorbonne Université UMR S1127, Institut du Cerveau et de la Moelle épinière (ICM), Social and Affective Neuroscience (SAN) Lab, Paris, France; Science & Technology of Music and Sound (STMS), UMR 9912 (CNRS/IRCAM/SU), Paris, France

## Abstract

In social interactions, people have to pay attention both to the *what* and *who*. In particular, expressive changes heard on speech signals have to be integrated with speaker identity, differentiating e.g. self- and other-produced signals. While previous research has shown that self-related visual information processing is facilitated compared to non-self stimuli, evidence in the auditory modality remains mixed. Here, we compared electroencephalography (EEG) responses to expressive changes in sequence of self- or other-produced speech sounds, using a mismatch negativity (MMN) passive oddball paradigm. Critically, to control for speaker differences, we used programmable acoustic transformations to create voice deviants which differed from standards in exactly the same manner, making EEG responses to such deviations comparable between sequences. Our results indicate that expressive changes on a stranger’s voice are highly prioritized in auditory processing compared to identical changes on the self-voice. Other-voice deviants generate earlier MMN onset responses and involve stronger cortical activations in a left motor and somatosensory network suggestive of an increased recruitment of resources for less internally predictable, and therefore perhaps more socially relevant, signals.

## 1 Introduction

In social interactions, people have to process continuous changes not only in the vocal and facial expressions of their interlocutors, but also in the feedback from their own facial and vocal expressions. There is a long-ranging debate in the social-cognitive and meta-cognitive communities (James, 1884; Frith, 2012) about the mechanistic primacy of both types of inputs: on the one hand, the social-cognitive interpretation of other agents is believed to mobilize simulation mechanisms which supplement the processing of exteroceptive input (Gallese et al, 2004; Niedenthal, 2007). On the other hand, vocal (Aucouturier et al, 2016) and facial feedback (Laird and Lacasse, 2014, but see also Wagenmakers et al, 2016) paradigms suggest that metacognitive evaluations of e.g. one’s own emotional state are influenced by proprioceptive inputs (the sound of our voice, the motor pattern of our face) that are processed *as if* they were external stimuli. In the voice domain in particular, the question remains whether there are fundamental mechanistic differences between e.g. hearing one’s own voice suddenly change its pitch to sound brighter and happier, and processing the exact same cues on the voice of a conversation partner.

Electrophysiological indices of self and other-stimulus processing have provided mixed evidence to this question. Various studies show converging evidence that self-related visual stimuli are prioritized in the brain (Apps and Tsakiris, 2014). For instance, images of the self-face elicit faster responses and recruit greater attentional resources than representations of another person (Pannese and Hirsch, 2011; Tacikowski and Nowicka, 2010; Sel et al, 2016). However, self-generated auditory stimuli do not necessarily show the same pattern: while participants’ own voice evoked larger N2 and P3 when compared to a stranger’s voice in an active detection task (Conde et al, 2015), Graux and colleagues found that participants’ own voice evoked a smaller P3a amplitude than the voice of a stranger or a familiar other when using passive oddball paradigms (Graux et al, 2013, 2015).

Critically, these previous studies have used designs in which self- and other-stimuli are alternated. While such a contrast sheds light on the relative saliency of self-voice deviants in a context of other-voices standards, it does not address the question of whether we process expressive changes in our own voice (i.e. self-voice deviants in a sequence of self-voice standards) in the same way as in the voice of others (i.e. other-voice deviants in a sequence of other-voice standards).

On the one hand, the processing of expressive changes in a sequence of self-voices may be facilitated because the self-voice is a familiar signal. Visual paradigms have consistently shown that deviants among familiar letters or shapes elicit faster mismatch responses (e.g. Sulykos et al, 2015), and similar results were found contrasting deviants in culturally familiar sounds (e.g. the Microsoft Windows chime) with deviants in sequences of the same sounds played backwards (Jacobsen et al, 2005). A self-voice advantage would also be consistent with results documenting facilitating effects of language- or speaker-familiarity on phonological and semantic processing (Chen et al, 2016; Fleming et al, 2014).

On the other hand, the processing of expressive cues in a sequence of other-voices may be facilitated because the other-voice is less predictable, more socially relevant, and thus warrants more/faster reorientation of attention than self-stimuli. There are known effects of social relevance on mismatch responses in the visual and auditory modalities, notably when manipulating the communicative nature of the signals: in sequences of emotional face stimuli, Campanella and colleagues (2002) found earlier and larger mismatch responses to changes of expressions that led to a different emotional appraisal (e.g. a happy face in a sequence of sad faces) than to a different depiction of the same emotion (see also Bayer et al, 2017; Kovarski et al, 2017). In the auditory domain, affiliative signals such as laughter evoke larger MMN than a non-affiliative growl (e.g. Pinheiro et al, 2017b), vowels expressing fear evoke both an earlier and larger MMN response than expressions of happiness and sadness (Carminati et al, 2018) and changes of the same intensity elicit larger MMNs on vocal than nonvocal stimuli (Schirmer et al, 2007), all of which can be interpreted as an effect of social relevance. Finally, not only the auditory stimulus itself, but also the context in which it is presented seems to affect preattentive change detection processes. In an oddball paradigm using both intensity and frequency deviants of pure tones, Pinheiro et al (2017a) reported a smaller MMN in response to deviants presented when participants looked at negative images compared to both positive and neutral images. Similarly, MMN responses to happy two-syllable deviants have shorter peak latencies when participants receive fear-reducing testosterone rather than placebo (Chen et al, 2014a). Even non-vocal tones modulated in F0 and F0 variation to match vocal expressions of affect are sufficient to evoke MMNs (Leitman et al, 2011).

One technical obstacle to comparing mismatch responses to expressive deviants in self and other-voice sequences, however, is the need to control for similar changes to occur in both contexts. When relying on participant voices, it is always possible that one speaker expresses a given emotional change more clearly or loudly than another speaker (Jürgens et al, 2015), or with different cues (e.g. louder vs higher pitch), such that any difference observed in processing such changes cannot be unambiguously attributed to self/other processing differences, rather than individual production differences.

To make such sequences amenable to an MMN paradigm, we used a novel voice-transformation software tool (DAVID, Rachman et al, 2018) in order to create voice deviants which, while being recognized as authentic emotional changes for both types of speaker, utilize *exactly* the same cues in *exactly* the same manner (e.g. a 50-cent pitch increase on the second syllable of the word) in both contexts. Previous studies using DAVID have demonstrated that transformed voices are perceived as natural emotional expressions, and create the same explicit and implicit emotional reactions as authentic emotional expressions (Aucouturier et al, 2016). In the present study, we used DAVID to apply identical expressive changes to both self- and other-voice stimuli, and used an event-related potential (ERP) mismatch negativity (MMN) paradigm to examine whether the processing of these controlled changes is affected by speaker identity.

## 2 Methods

### 2.1 Participants

Twenty-five healthy, right-handed female participants took part in this study (27 came in for voice recordings, but two were not able to do the EEG session), two of which were excluded from analysis due to excessive EEG artifacts in the EEG, leaving 23 participants in the final analysis (mean age=21.2, SD=1.8 years).

An additional twenty right-handed female participants took part in a follow-up behavioral study comprising an emotion categorization task. One participant was excluded because of missing data due to technical problems, leaving 19 participants in the final analysis (mean age=21.4, SD=2.1 years). Participants in this second group did not partake in the EEG experiment.

For both studies, we selected only female participants because the voice transformations we used worked more reliably for female than for deep, lower-pitch male voices (Rachman et al, 2018). The experimental protocol was approved by INSEAD’s institutional review board and all participants gave written informed consent before the start of the study. Participants reported normal or corrected to normal vision, normal hearing, and an absence of neurological or psychiatric illness. They were financially compensated for their participation.

### 2.2 Stimuli

Participants came to the lab one week prior to the EEG experiment for a voice recording session. The recordings took place in a sound-attenuated booth, using a headset microphone (DPA d:fine 4066), an external sound card (RME UCX Fireface), and Garage-Band software (Apple Inc.) with a 44.1 kHz sampling rate and 16-bit resolution. Participants were asked to read a list of twenty disyllabic neutral words and six disyllabic pseudo-words with a neutral intonation (Table S1). All sounds were normalized at 70 dBA using a Matlab toolbox (Pampalk, 2004). Because only the recordings of the pseudoword */ba-ba/* were used during the EEG session, these sound files were also normalized in time to have a duration of 550 ms using superVP/audiosculpt software. To ensure comparable amounts of vocal diversity in ‘self’ and ‘other’ stimuli, participants were grouped in pairs such that the ‘self’ voice (SV) of one participant served as the ‘other’ voice (OV) for the other participant, and vice-versa.

Finally, we processed all recordings with the DAVID software platform (Rachman et al, 2018) to generate happy and sad deviants from the standard utterance, by combining audio effects such as pitch shift (increasing the standard’s pitch by 50 cents in the happy deviant, and decreasing by 70 cents in the sad deviant), inflection (increasing the beginning of the second syllable by an extra 70 cents in happy) and filtering (increasing high-frequency energy with a high-shelf filter in happy, and decreasing high-frequency energy with a low-shelf filter in sad; see Table 1 for parameter values). Crucially, using such programmable emotional transformations ensured that, in both the self and other sequences, deviants differed from the standards in exactly the same manner, making EEG responses to such deviations comparable between sequences (Figure 1).

**Table 1:** Deviant parameter values

|  | Happy | Sad |
| --- | --- | --- |
| **Pitch** |  |  |
| shift, *cents* | +50 | -70 |
| **Inflection** |  |  |
| duration, *ms* | 500 | – |
| min., *cents* | -200 | – |
| max., *cents* | +140 | – |
| **Shelf filter** |  |  |
| cut-off, *Hz* | >8000 | <8000 |
| slope, *dB/octave* | +9.5 | -12 |

Parameter values of the happy and sad transformations used in this study (refer to Rachman et al (2018) for details).

**Figure 1:**
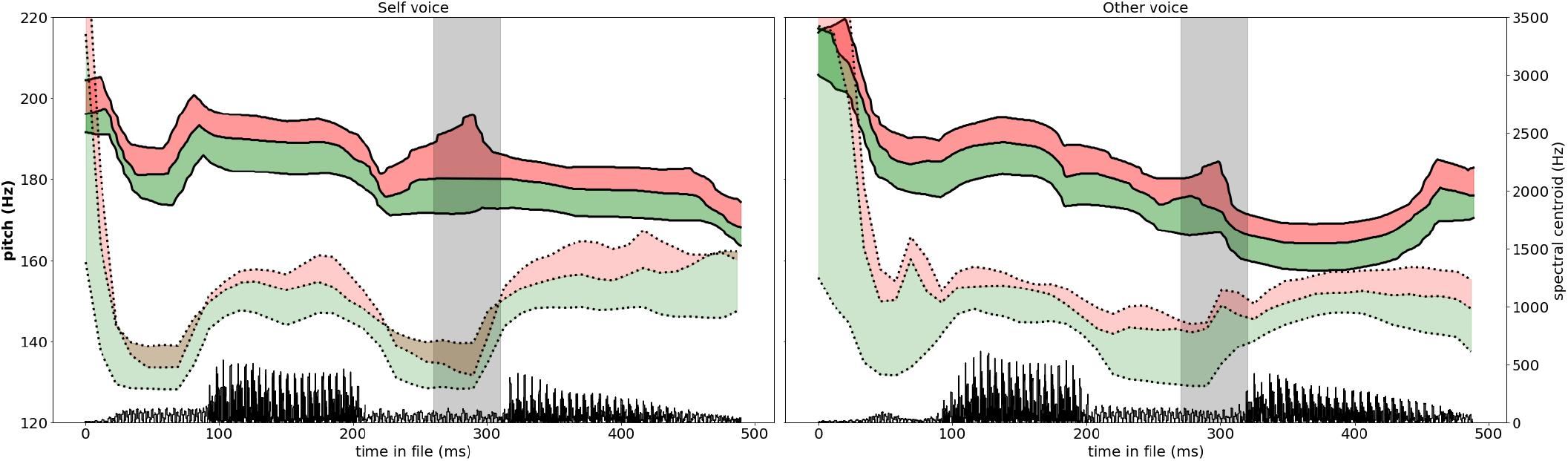
Acoustic content of two representative stimuli used in the MMN experiment. Solid line, black: pitch of the standard; red: increase of pitch in the happy deviant; green: decrease of pitch in the sad deviant. Shaded area indicates second-syllable inflection in the happy deviant. Dotted line, black: spectral centroid (centre of mass) of the standard; red: high-frequency energy added in the happy deviant; green: high-frequency energy removed in the sad deviant. Bottom, black: half-corrected waveforms of the standard. Left: participant’s own voice (SELF). Right: another participant’s voice (OTHER).

### 2.3 Oddball paradigm

We used an oddball paradigm with two different sequences: one ‘self sequence’ and one ‘other sequence’. In the ‘self sequence’, the neutral recording of the SV served as the standard stimulus and the ‘happy’ and ‘sad’ transformation of the standard stimulus served as the two emotional deviants. Following the same logic, the ‘other-sequence’ used the neutral recording of the OV as the standard and its ‘happy’ and ‘sad’ transformations as deviants (see Fig. 2). Additionally, both sequences also contained an identity deviant to try to replicate previous studies by Graux and colleagues (2013; 2015): the neutral SV was presented as the identity deviant in the ‘other sequence’ and vice versa (see section Replication of Graux et al. (2015) for further information). We counterbalanced the order of the sequences across participants. Each sequence contained 1080 stimuli in total with the standard stimulus occurring 80% of the time and each of the three deviant stimuli (‘happy’, ‘sad’ and ‘identity’) occurring 6.7% of the time (72 stimuli). Each sequence started with 10 standard stimuli and 2-7 standards occurred between successive deviants. All stimuli lasted 550 ms and were presented with a stimulus onset asynchrony (SOA) of 1000 ms.

**Figure 2:**
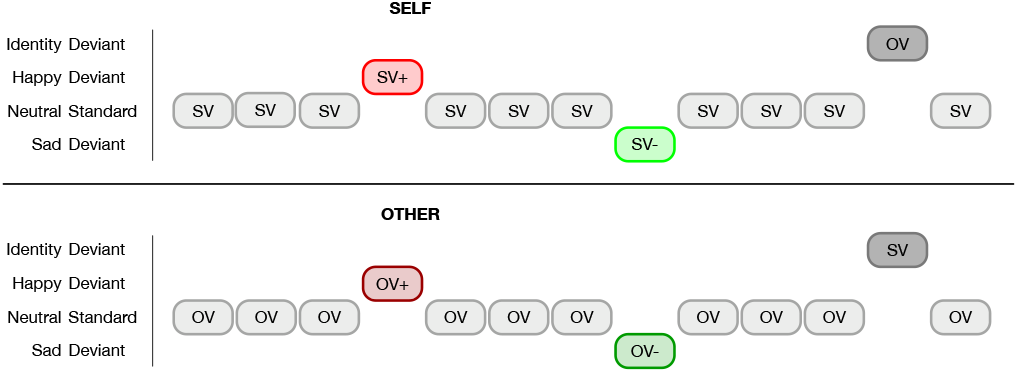
Schematic representation of the oddball sequences for the self (above) and other (below) conditions. In the self-voice sequences, standards are neutral self-voice (SV) and deviants are happy (SV+) and sad (SV-) manipulations of the standard, as well as one other-voice recording of the same word (OV). In the other-voice sequences, standards are neutral other-voices (OV) and deviants are happy (OV+) and sad (OV-) manipulations as well as one self-voice recording of the same word (SV).

### 2.4 Discrimination and intensity rating tasks

To test whether participants were able to distinguish their own voice from a stranger’s voice, they performed a behavioral self-other discrimination task. Five bisyllabic words and three pseudowords (Table S1), produced by the participant and another person (the same ‘other’ as was presented during the EEG recording) were presented in neutral, happy, and sad versions. Participants were asked to indicate for each stimulus if it was their own voice or the voice of someone else. For the intensity task, participants listened to the same stimuli as in the discrimination task and were asked to rate the emotional intensity of the voice on a continuous rating scale (0-100). All stimuli were separately randomized for both tasks.

### 2.5 Follow-up emotion recognition task

To test whether the emotional transformations were correctly recognized, a second group of N=20 female participants performed an emotion categorization task with the same stimuli as above (five bisyllabic and three pseudo words). Participants were presented with pairs of the same word produced by the same speaker (SV or OV). The first stimulus was always an original, non-manipulated recording and the second stimulus was either a neutral recording or transformed using the ‘happy’ or ‘sad’ effect. Participants were then asked to categorize the second stimulus in a 3-category emotion recognition task (neutral-happy-sad).

### 2.6 Procedure

During the EEG recordings, subjects were seated in front of a computer screen (55*×*32 cm) on which they watched a silent subtitled movie. Participants were asked to pay attention to the movie and to ignore the sounds. Auditory stimulus presentation was controlled with PsychoPy (Peirce, 2007) and sounds were delivered through Sennheiser CX 300-II earphones at 70 dB SPL.

### 2.7 EEG data acquisition

Electroencephalographic (EEG) data were recorded from 63 scalp locations (actiCHamp, Brain Products GmbH, Germany) with a sampling rate of 500 Hz, relative to a nose tip reference, and filtered with a bandpass of 0.01-100 Hz (12 db/octave roll-off). Four electrodes were placed on the left and right temples (horizontal electrooculogram [EOG]) and above and below the left eye (vertical EOG) to monitor eye movements and blinks respectively.

Pre-processing and statistical analyses were performed in FieldTrip (Oostenveld et al, 2011). Offline, the continuous data were re-referenced to the average of the left and right mastoid electrodes (TP9 and TP10) and filtered with a 0.1 Hz high-pass filter (Butterworth, 12 dB/octave roll-off) and a 30 Hz low-pass filter (Butterworth, 48 dB/octave roll-off). The data were then visually inspected to remove epochs with artifacts, such as muscle activity and signal drifts. Next, eye blinks and movements were corrected using the fast independent component analysis (fastICA) method.

To get a better estimation of the MMN, we equated the number of deviants and standards by randomly selecting 69 standards (as many as the mean number of deviants after artifact rejection) that immediately preceded a deviant in the self and other sequences. Individual EEG epochs were averaged separately for each type of standard stimulus (self, other; Fig. 3) and deviant stimulus (neutral self, neutral other, happy self, happy other, sad self, sad other), with a 200 ms pre-stimulus baseline and a 700 ms post-stimulus period. After artifact rejection, each subject had at least 75% trials remaining in each condition and the number of trials did not differ across conditions (Self standard: M=831.5, happy: M=69.7, sad: M=69.4; Other standard: M=831.3, happy: M=68.7, sad: M=69.0; *ps* > .05). Finally, four difference waves were calculated by subtracting the grand average waveform of the standard stimuli from each of the deviant grand averages within each sequence type (i.e., for each speaker separately), yielding the following conditions: ‘Happy Self’, ‘Happy Other’, ‘Sad Self’, ‘Sad Other’ (Fig. 4).

**Figure 3:**
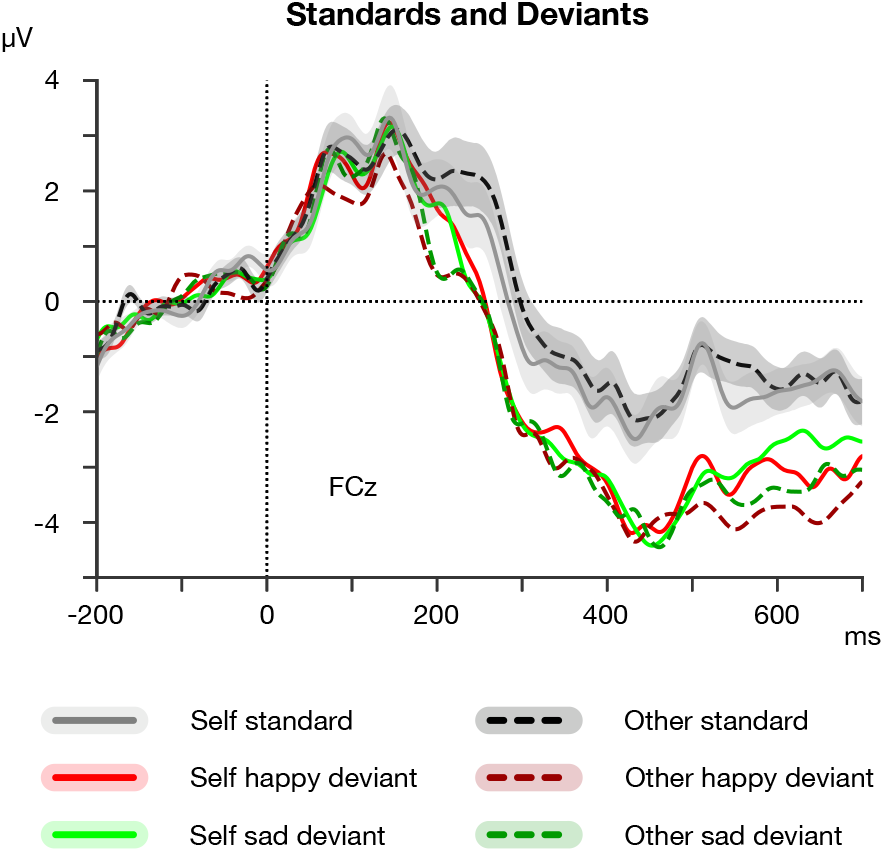
Grand-average ERPs to the self (solid lines) and other (dashed lines) standard and deviant stimuli. Shaded area represents bootstrap standard error of the mean (SEM).

**Figure 4:**
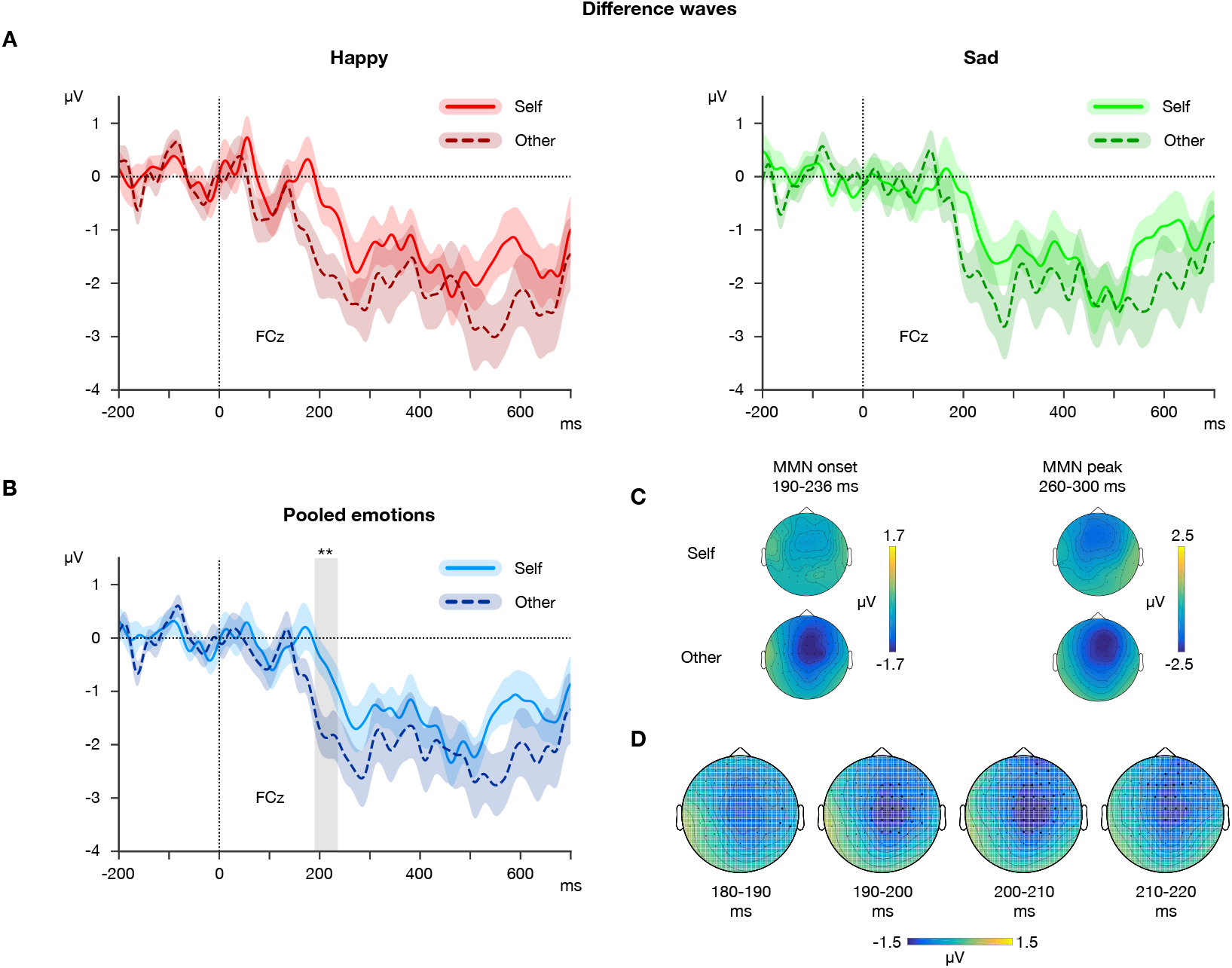
(A) Differences waves of the happy (red) and sad (green) transformations on the self (light) and other (dark) voice. (B) Differences waves of the pooled happy and sad deviants of the self (light) and other (dark) voice. Shaded area represents bootstrap SEM, \*\**p* < .01. (C) Topographies of the pooled happy and sad deviants of the self and other voice at MMN onset and peak. (D) Significant cluster of the contrast between the ‘other’ and ‘self’ difference waves represented in four 10 ms time windows between 180 and 220 ms. Highlighted channels belong to the cluster and were significant across the whole time 10 ms window.

### 2.8 Statistical analyses

Statistical analyses were conducted in Python 2.7. The alpha level was set at .05 and all statistical tests were two-tailed.

#### 2.8.1 Behavioral data

Accuracy scores and ratings were computed from the discrimination and intensity rating tasks respectively. We conducted one-sample *t*-tests on the accuracy scores to test whether SV and OV were discriminated above chance level (50%). For the intensity task, we conducted a two-way repeated-measures analysis of variance (rmANOVA) with the factors Emotion (neutral, happy, sad) and Identity (self, other). Significant main effects and interactions were followed up with Tukey HSD for post-hoc comparisons.

#### 2.8.2 EEG data

EEG data were analyzed using cluster-based statistics implemented in FieldTrip (Maris and Oostenveld, 2007). In total, four cluster-based permutation tests were performed: one on the standard grand-averages to test for an effect of speaker identity and three on the difference waves to investigate main effects of identity and emotion and their interaction. For the interaction we first calculated the difference between the happy and sad difference waves for each speaker identity separately before entering these data into the analysis. Based on prior hypotheses about the temporal location of the MMN component (e.g. Graux et al, 2013; Pinheiro et al, 2017b; Beauchemin et al, 2006), analyses were carried out within a 50-300ms time window across all electrodes. For each cluster-based permutation test we first conducted pairwise *t*-tests between two conditions at each time-point and channel in the predefined time-window and region of interest. The cluster-level statistic was defined as the sum of all individual *t*-statistics that belong to the cluster. This procedure then controls the type I error rate by evaluating the cluster-level statistic under the randomization null distribution of the maximum cluster-level test statistic, obtained by 5000 permutations.

As an alternative parametric analysis strategy, we analyzed the mean MMN amplitude and MMN peak and onset latencies within a region of interest comprising electrodes F1, Fz, F2, FC1, FCz, FC2, C1, Cz, and C2. We extracted the mean amplitude over a 40 ms time window around the averaged MMN peak across conditions, participants, and electrodes (280 *±* 20 ms), to avoid a possible bias introduced by differences in conditions. We extracted the MMN onset and peak latencies using a jackknife procedure and tested for differences in the four conditions (Identity *×* Emotion). The jackknife procedure improves statistical power by taking the latencies of the grand average using a leave-one-out method (Kiesel et al, 2008; Ulrich and Miller, 2001): for N=23 participants, we calculated 23 grand averages, each leaving out one of the participants and including the other 22. We then determined the onset latency for each of these 23 grand averages as the time where the difference wave reached 50% of the MMN peak amplitude. In a similar way, we defined the MMN peak latency as the time at which the difference wave reached the most negative amplitude. These values were entered into two separate two-way rmANOVAs with identity (Self, Other) emotion (Happy, Sad), antero-posterior site (Frontal, Frontocentral, Central) and lateralization (1-line, z-line, 2-line) as within-subject factors. Finally, we divided the resulting *F*-value by (*N−*1)^2^ to correct for the artificially low error variance introduced by the leave-one-out procedure (Ulrich and Miller, 2001). Furthermore, Greenhouse-Geisser correction for non-sphericity was applied when necessary. We report uncorrected degrees of freedom and corrected *p*-values.

#### 2.8.3 Source localization

Estimation of cortical current source density was performed with Brainstorm (Tadel et al, 2011). The cortical current source density mapping was obtained from a distributed source model of 15000 current dipoles. The dipoles were unconstrained to the cortical mantle of a generic brain model built from the standard MNI template brain provided in Brainstorm. EEG electrode positions were determined for each subject using a CapTrak system (Brain Products GmbH, Germany) and aligned to the standard MNI template brain. The forward model was computed with the OpenMEEG Boundary Element Method (Gramfort et al, 2010). A noise-covariance matrix was computed for each subject by taking the 200 ms baseline period of each trial and was taken into account in the inversion algorithm. The cortical current source density mapping was then obtained for each subject from the time series of each condition by means of the weighted minimum-norm estimate (wMNE). Z-scored cortical maps across all conditions were used to define the regions of interest that are activated irrespective of emotion and identity within the time window in which there was a significant difference between self and other conditions. Regions of interest (ROIs) contained at least 30 vertices with a z-score above 60% of the maximum z-score. To analyze the cortical sources of the difference waves we performed paired *t*-tests for each vertex within the defined ROIs, taking the mean values across the 190-230 ms window. This time window was chosen to span the interval between the average MMN onset latency in the OV condition (190 ms) and the average MMN onset latency in the SV condition (236 ms), in order to identify sources for the activity explaining the effect (see Results section). Activations within an ROI were considered significant whenever at least ten adjacent vertices reached statistical significance.

#### 2.9 Replication of Graux et al. (2015)

In addition to the above procedure, we included extra stimuli to replicate the identity mismatch response of Graux et al (2015), namely one neutral-other deviant (P=.067) in the ‘self’ sequences (the same stimulus that served as standard in the ‘other’ sequence), and one neutral-self deviant (P=.067) in the ‘other’ sequences (the same stimulus that served as standard in the ‘self’ sequence). Difference waves were calculated by subtracting (neutral) standards that immediately preceded the identity deviant of one sequence (e.g. ‘other’ standard from ‘other’ sequence) from the neutral deviant of the same identity in the other sequence (e.g. ‘other’ deviant from ‘self’ sequence). While not statistically significant, the pattern of responses to both types of deviants was consistent with Graux et al (2015), with larger P3a for ‘other’ deviants than ‘self’ (see Fig. S1). These results are not further discussed in this paper.

## 3 Results

### 3.1 Behavioral results

In the post-EEG task, participants were tested on a variety of self- and other-voice stimuli, processed or unprocessed with emotional manipulations, and asked to evaluate both whether these were examples of the self-voice, and how emotionally intense the samples were.

The accuracy of self-other discrimination was greater in OVs than in SVs (main effect of speaker identity: *F* (1, 22) = 80.7, *p* < .001), which is easily explained by the fact that it is easier to misattribute sounds from the self to (an infinite possibility of) other identities, than the other way around. There was also a main effect of emotional manipulation on discrimination accuracy (*F* (2, 44) = 19.2, *p* < .001) and an identity *×* emotion interaction (*F* (2, 44) = 20.7, *p* < .001), showing that manipulated self-voices were more easily confused for other identities than non-manipulated voices. Self-other discrimination was more accurate than chance for both neutral and emotional other-voices (*t*s(22) *>* 19, *ps* < .001), more accurate than chance in the neutral (*t*(22) = 9.66, *p* < .001) and sad self-voices (*t*(22) = 2.60, *p* < .05), but not in happy self-voices (*t*(22) = *−*1.07, *p* > .05; Fig. 5, Left).

**Figure 5:**
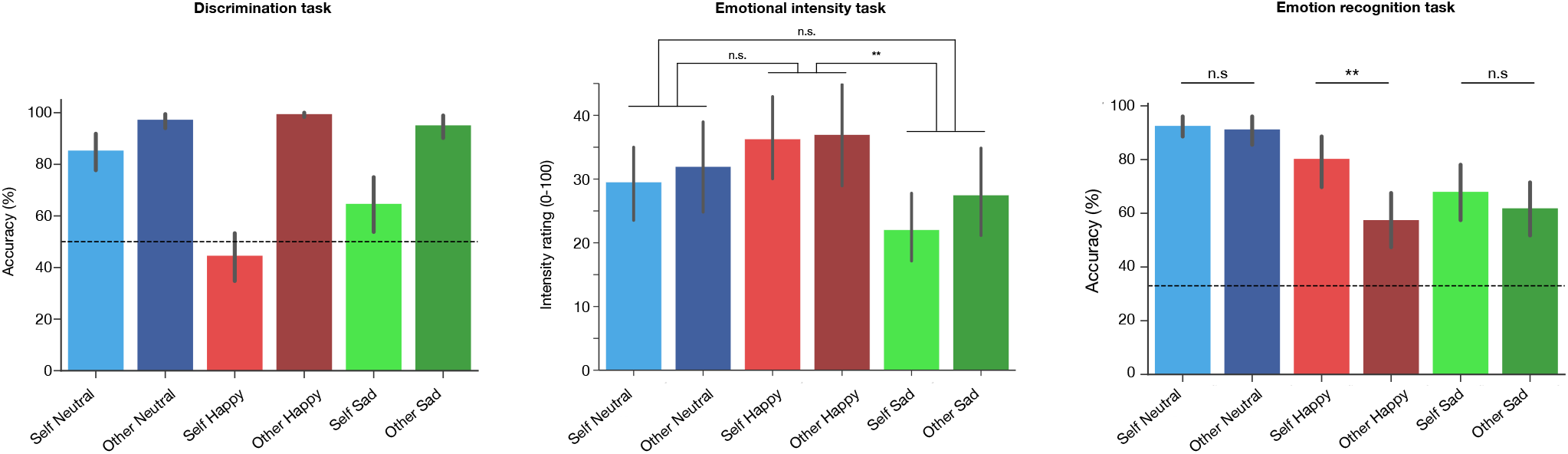
Left: Discrimination accuracy (%) for neutral, happy, and sad versions of SV and OV. Dotted line indicates chance level performance (50%). Middle: Intensity ratings (0-100) for neutral, happy, and sad versions of self and other voice. Right: Emotion recognition accuracy (%) for neutral, happy, and sad versions of SV and OV. Dotted line indicates chance level performance (33.3%). Error bars represent standard error of the mean. \*\**p* < .01

As expected, ratings of emotional intensity showed a main effect of emotion (*F* (1, 22) = 21.63, *p* < .001). Sad voices were rated as less intense than happy (*p* < .01), but neither emotion differed from neutral (see Fig. 5, Middle). There was no effect or interaction with speaker identity, confirming that emotional deviants were comparable in both types of sequences.

In the additional emotion recognition task, accuracy was greater in SV than in OV (main effect of speaker identity: *F* (1, 18) = 15.14, *p* < .01). We also found a main effect of emotion (*F* (2, 36) = 20.59, *p* < .001), as well as an emotion *×* identity interaction (*F* (2, 36) = 5.31, *p* < .05). Follow-up paired sample *t*-tests revealed that the happy transformation was better recognized on the SV than on the OV: *t*(18) = 4.03, *p* < .01 (Bonferroni correction; Fig. 5, Right).

### 3.2 Standards

The cluster-based permutation test and rmANOVAs did not reveal any differences between the self and other standard conditions (*ps* > .05).

### 3.3 Difference waves

Difference waves showed a relatively small (−2*µ*V) fronto-central negativity peaking at 280 ms*±*20ms, compatible with a MMN (Figure 4). The difference waves were also re-referenced to the nose reference to ensure the typical polarity inversion between Fz/Cz and the mastoid electrodes. However, because mastoid-referenced averages typically show a better signal-to-noise ratio than the nose-referenced averages, the former were used in all subsequent analyses (Kujala et al, 2007; Martínez-Montes et al, 2013).

We found a significant cluster when testing for a main effect of identity (Monte Carlo *p* < .05; Fig. 4D) but none for a main effect of emotion or an interaction. Parametric analyses with the jackknife procedure revealed that this difference was driven by the onset of the MMN, rather than its peak. There was a main effect of identity on the MMN onset latency (*F*_corrected_(1,22) = 10.14, *p* < .01), with the OV onset latency at 190 ms, compared to 236 ms in the SV condition, a considerable difference of 46 ms (see Fig. 4A-C for the difference waves and topographies). There were no effects of emotion, electrode antero-posterior location or lateralization on onset latency, nor was there a significant interaction between any of the factors. In contrast, no main effects of identity or emotion were observed in the amplitude (−2 *µ*V) or the peak latency (280*±*20 ms) of MMN peak and no interaction effects were observed on the MMN peak latency. The repeated-measures ANOVA on the mean MMN amplitude showed only an identity *×* lateralization interaction effect (*F* (2, 44) = 7.04, *p* < .01), but follow-up analyses at each antero-posterior site (Frontal, Fronto-central, Central) did not reveal an effect of identity (all *ps* > .05).

### 3.4 Sources

ROIs identified using source activation maps across all conditions in the 190-230 ms window (spanning the difference between other- and self-MMN onset latencies) included bilateral regions in the precentral gyri, large insulo-temporal regions in the right hemisphere and large fronto-parietal regions in the left hemisphere. Source activations for OV versus SV in these ROIs were stronger in the left precentral gyrus/sulcus (47 vertices), and the left postcentral gyrus (16 vertices). (Fig. 6).

**Figure 6:**
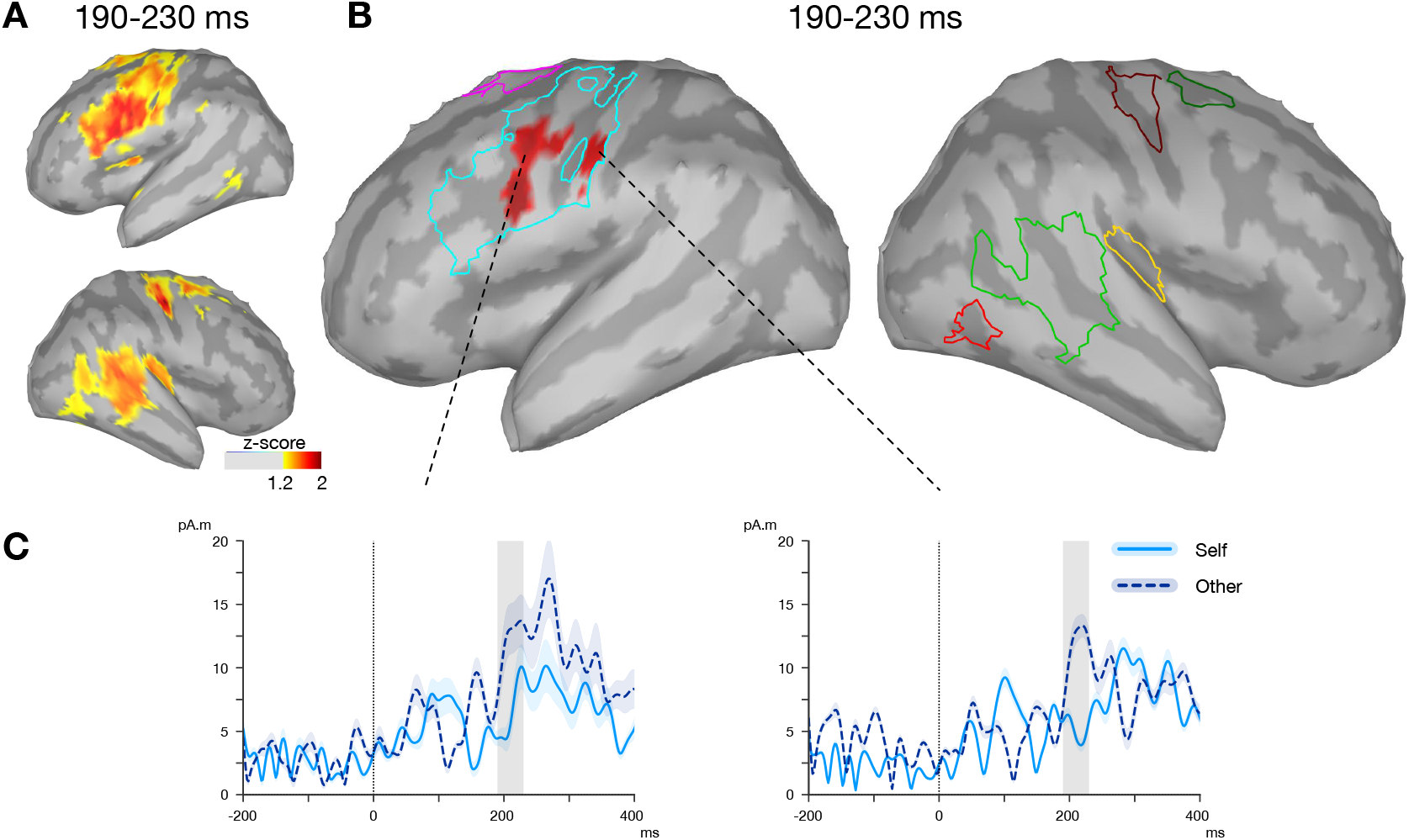
(A) Source localizations across all conditions in the 190-230 ms showing maxima of activation (>60%, z-scores) to determine Regions of Interest (ROIs). (B) Modulations of cortical activity as a function of speaker identity in the 190-230 ms time window. Only clusters containing at least 10 contiguous vertices with *p* < .05 in this time window were considered statistically significant. The source activations are color-coded only for *t*-values corresponding to *p* < .05. (C) Time courses of the grand mean amplitude of the current sources in each activated region for self and other conditions. Shaded areas represent the standard deviation, the grey area represents the 190-230 ms time window in which the analyses took place.

## 4 Discussion

Changes in vocal cues can communicate a person’s social attitudes or emotional states and are thus important to process in social interactions. The present study investigated whether the same emotion-related acoustic changes (pitch variations, inflections, and timbre) are processed differently on the self-voice compared to the voice of a stranger.

### 4.1 Behavior

Self/other discrimination rates for transformed versions of the SVs were lower than for the neutral SVs, which suggests that our emotional manipulations, and notably the happy effect, affected identity perception to a certain extent. While it is difficult to relate such subsequent, explicit recognition scores to the implicit processes occurring during the earlier oddball procedure (see e.g. Candini et al, 2014), it remains possible that some of the participants processed deviants in self-voice sequences as differing both in emotional tone and speaker identity. However, it appears implausible that such misattributed deviants in SV sequences should drive the greater MMN onset latencies seen in these sequences, compared to OV sequences. First, deviants misattributed as OVs in sequences of SVs have been traditionally associated with greater, rather than lower saliency (e.g. greater P3a amplitude in Graux et al, 2013), such that misattributions of identity in SV sequences should reduce, rather than accentuate the effect found here. Second, while behavioral data shows speaker identity is affected to a larger extent by the happy effect than the sad effect, EEG responses to happy and sad deviants did not differ.

In other auditory tasks, a sound’s increased emotional or social relevance often creates perceptual biases that make them appear louder or more intense (Neuhoff, 1998; Asutay and Västfjäll, 2012). Here, self and other-voice emotions did not differ in their perceived emotional intensity. However, while manipulated emotions were recognized well above chance level for both self and other voices, happy (and to a lesser extent, sad) manipulations were better recognized when participants heard them on their own voice, rather than that of an unfamiliar stranger. This pattern of results is in line with a number of studies showing better recognition or prediction accuracy when one observes one’s own actions than when one observes another person’s actions (Knoblich and Flach, 2001; Tye-Murray et al, 2015), and can also be explained by better sensory resolution for the familiar sounds of one’s own voice, similarly perhaps to the language familiarity effects seen with native vs foreign-language speaker discrimination (Fleming et al, 2014).

### 4.2 MMN onset latency

Across all conditions, the MMN peak latency was a relatively late 280 ms. MMN usually peaks at 150-250 ms from change onset, with this peak latency getting larger with the decreased magnitude, or increased processing difficulty, of stimulus change (Garrido et al, 2009). It is possible that the late peak latency observed here reflects a late onset of observable stimulus change in our two-syllable words. In particular, spectral changes associated with happy or sad deviants may only become manifest on the vowel portion of the first syllable (onset ca. 100 ms, see Figure 1). In similar studies of two-syllable emotional words with a variety of changes (e.g. consonant duration, omission of second syllable, etc.), Pakarinen et al (2014) report MMN peak latencies ranging between 126-355 ms post stimulus-onset, and Chen et al (2014b) a peak MMNm of 265 ms; in contrast, with single-vowel stimuli involving more immediate timbre changes and no initial consonant, Carminati et al (2018) report MMN latencies around 200 ms. Future work should better document the temporal profile of physical information available in the signal to discriminate deviants from standards, in order to more precisely determine the chronometry of their auditory processing.

We observed no difference in MMN amplitude and peak latency, but an earlier MMN onset for emotional deviants on the OV compared to the SV. This MMN onset latency effect was seen in both emotional transformations, and amounted to a considerable difference of 46 ms. Because we did not find any significant difference between the self and other conditions on the waveform of the standard stimuli, and because both self and other deviants were generated from the standards with identical algorithmic procedures, it is unlikely that such a large onset effect results from the differential processing of the standards, or differences in refractory states (Jacobsen and Schröger, 2001).

The shorter MMN onset latency in the OV condition rather suggests that changes on a stranger’s voice are highly prioritized in auditory processing. This is in contrast with the increased saliency of self-stimuli in the visual domain (Apps and Tsakiris, 2014; Sel et al, 2016), but consistent with the idea that other-stimuli are more socially relevant (Pinheiro et al, 2017b; Schirmer et al, 2007). In a recent study, effects of emotion were seen earlier in a communicative context when compared to a non-communicative context (Rohr and Abdel Rahman, 2015). It therefore appears possible that our design of other-deviants in a sequence of other-standards is implicitly treated as a context akin to social communication (“other speaking to self”), more so than changes embedded in a sequence of self-sounds.

It should be emphasized that only female participants were included in this study. While both women and men typically show an MMN response to emotional deviants, previous work has showed that this preattentive response can be amplified in women, possibly because of a greater social relevance of emotional information for women (Schirmer et al, 2005). Importantly, this amplification seems to be specific to vocal sounds and has not been found in nonvocal sounds (Hung and Cheng, 2014). As such, it remains to be determined whether male participants show a similar difference in MMN onset latency as what we report here.

### 4.3 Source activations

Source estimations during the MMN onset temporal window (190-230 ms) across all conditions showed activations in the right insulo-temporal region and the left fronto-parietal region. Right-lateralized temporal activations are in line with previous MMN studies that reported right activations for pitch deviants in tones and voice (Jiang et al, 2014; Lappe et al, 2016). In addition, the right anterior insula is involved in processing vocal emotions Belin et al (2004), and has also been associated to MMN responses to emotional syllable deviants (Chen et al, 2014b).

The interpretation of EEG source analysis should remain conservative. Here, activity discriminative of self and other mismatches did not occur within the typical supra-temporal or frontal MMN generators (Garrido et al, 2009), which suggests that processing other-voice stimuli was accompanied neither by any detectable enhancement of sensory processes, nor by any switch of attention. Neither did activity discriminative of self and other occur within the predominantly right-lateralized regions previously associated with speaker identity tasks, such as the right temporoparietal junction (Schall et al, 2015) and right inferior frontal gyrus (Kaplan et al, 2008), or with MMN sources associated with emotional vocal stimuli such as the right anterior insula (Chen et al, 2014b). Instead, when contrasting responses to ‘self’ and ‘other’ deviants within the above ROIs, we found increased activations in the left precentral gyrus/sulcus and the left postcentral gyrus for deviants on the other voice.

These regions suggest that emotional vocal deviants recruit a network of motor and somatosensory areas which are increasingly thought to be involved in mapping heard speech onto articulatory representations (Scott and Johnsrude, 2003; Evans and Davis, 2015; Skipper et al, 2017). The left somatomotor cortex in particular has been associated with phoneme discrimination tasks (Sato et al, 2009), and appears to be especially recruited for more effortful conditions involving noisy (Hervais-Adelman et al, 2012; D’Ausilio et al, 2012) or non-native speech (Wilson and Iacoboni, 2006), in which articulatory representations may provide a processing advantage. In the visual modality, left somatosensory areas have also been associated with unpredicted deviations from the self face (Sel et al, 2016), or facial emotion recognition in the other (Sel et al, 2014), both of which are also believed to involve processes of embodied simulation or predictions. Earlier activity in this network of regions for the other-voice deviants is therefore compatible with a greater recruitment of resources for less-internally predictable signals such as speech produced by an unfamiliar stranger, for which listeners may lack an adequate internal template - a fact that can also explain that recognizing emotions in a separate explicit task was also more difficult on non-self voices.

In sum, expressive changes on a stranger’s voice are highly prioritized in perceptual processing compared to identical changes on the self-voice. Other-voice deviants generate earlier MMN responses and involve activity in a left motor/somatosensory network suggestive of greater recruitment of resources for less internally predictable, and therefore perhaps more socially relevant, signals.

## 5 Funding

This study was supported by ERC Grant StG 335536 CREAM to JJA.

### 6 Acknowledgements

All data were collected at the Centre Multidisciplinaire des Sciences Comportementales Sorbonne-Université-INSEAD. The authors thank Maël Garnotel for his help with the data collection, Nathalie George for advice on the source analyses, and Marie Gomot for comments on the manuscript.

## References

Apps MA, Tsakiris M (2014) The free-energy self: a predictive coding account of self-recognition. Neuroscience & Biobehavioral Reviews 41:85–97

Asutay E, Västfjäll D (2012) Perception of loudness is influenced by emotion. PloS one 7(6):e38,660

Aucouturier JJ, Johansson P, Hall L, Segnini R, Mercadié L, Watanabe K (2016) Covert digital manipulation of vocal emotion alter speakers’ emotional states in a congruent direction. Proceedings of the National Academy of Sciences 113(4):948–953

Bayer M, Ruthmann K, Schacht A (2017) The impact of personal relevance on emotion processing: Evidence from event-related potentials and pupillary responses. Social Cognitive and Affective Neuroscience 12:1470–1479

Beauchemin M, De Beaumont L, Vannasing P, Turcotte A, Arcand C, Belin P, Lassonde M (2006) Electrophysiological markers of voice familiarity. European Journal of Neuroscience 23(April):3081–3086

Belin P, Fecteau S, Bedard C (2004) Thinking the voice: neural correlates of voice perception. Trends in cognitive sciences 8(3):129–135

Campanella S, Gaspard C, Debatisse D, Bruyer R, Crommelinck M, Guerit JM (2002) Discrimination of emotional facial expressions in a visual oddball task: an erp study. Biological psychology 59(3):171–186

Candini M, Zamagni E, Nuzzo A, Ruotolo F, Iachini T, Frassinetti F (2014) Who is speaking? Implicit and explicit self and other voice recognition. Brain and Cognition 92:112–117

Carminati M, Fiori-Duharcourt N, Isel F (2018) Neurophysiological differentiation between preattentive and attentive processing of emotional expressions on french vowels. Biological psychology 132:55–63

Chen B, Kitaoka N, Takeda K (2016) Impact of acoustic similarity on efficiency of verbal information transmission via subtle prosodic cues. EURASIP Journal on Audio, Speech, and Music Processing 2016(1):19

Chen C, Chen CY, Yang CY, Lin CH, Cheng Y (2014a) Testosterone modulates preattentive sensory processing and involuntary attention switches to emotional voices. Journal of neurophysiology 113(6):1842–1849

Chen C, Lee YH, Cheng Y (2014b) Anterior insular cortex activity to emotional salience of voices in a passive oddball paradigm. Frontiers in human neuroscience 8:743

Conde T, Goncalves OF, Pinheiro AP (2015) Paying attention to my voice or yours: An erp study with words. Biological psychology 111:40–52

D’Ausilio A, Bufalari I, Salmas P, Fadiga L (2012) The role of the motor system in discriminating normal and degraded speech sounds. Cortex 48(7):882–887

Evans S, Davis MH (2015) Hierarchical organization of auditory and motor representations in speech perception: Evidence from searchlight similarity analysis. Cerebral Cortex 25(12):4772–4788

Fleming D, Giordano BL, Caldara R, Belin P (2014) A language-familiarity effect for speaker discrimination without comprehension. Proceedings of the National Academy of Sciences 111(38):13,795–13,798

Frith CD (2012) The role of metacognition in human social interactions. Philosophical Transactions of the Royal Society B: Biological Sciences 367:2213–2223, DOI 10.1098/rstb.2012.0123

Gallese V, Keysers C, Rizzolatti G (2004) A unifying view of the basis of social cognition. Trends in cognitive sciences 8(9):396–403

Garrido MI, Kilner JM, Stephan KE, Friston KJ (2009) The mismatch negativity: a review of underlying mechanisms. Clinical neurophysiology 120(3):453–463

Gramfort A, Papadopoulo T, Olivi E, Clerc M (2010) Openmeeg: opensource software for quasistatic bioelectromagnetics. Biomedical engineering online 9(1):45

Graux J, Gomot M, Roux S, Bonnet-Brilhault F, Camus V, Bruneau N (2013) My voice or yours? an electrophysiological study. Brain topography 26(1):72–82

Graux J, Gomot M, Roux S, Bonnet-Brilhault F, Bruneau N (2015) Is my voice just a familiar voice? an electrophysiological study. Social cognitive and affective neuroscience 10(1):101–105

Hervais-Adelman AG, Carlyon RP, Johnsrude IS, Davis MH (2012) Brain regions recruited for the effortful comprehension of noise-vocoded words. Language and Cognitive Processes 27(7-8):1145–1166

Hung AY, Cheng Y (2014) Sex differences in preattentive perception of emotional voices and acoustic attributes. Neuroreport 25(7):464–469

Jacobsen T, Schröger E (2001) Is there pre-attentive memory-based comparison of pitch? Psychophysiology 38:723–727

Jacobsen T, Schröger E, Winkler I, Horváth J (2005) Familiarity affects the processing of task-irrelevant auditory deviance. Journal of Cognitive Neuroscience 17(11):1704–1713

James W (1884) What is an emotion? Mind 9(34):188–205

Jiang A, Yang J, Yang Y (2014) Mmn responses during implicit processing of changes in emotional prosody: an erp study using chinese pseudo-syllables. Cognitive neurodynamics 8(6):499–508

Jürgens R, Grass A, Drolet M, Fischer J (2015) Effect of acting experience on emotion expression and recognition in voice: Non-actors provide better stimuli than expected. Journal of nonverbal behavior 39(3):195–214

Kaplan JT, Aziz-Zadeh L, Uddin LQ, Iacoboni M (2008) The self across the senses: An fMRI study of self-face and self-voice recognition. Social Cognitive and Affective Neuroscience 3:218–223

Kiesel A, Miller J, Jolicœur P, Brisson B (2008) Measurement of erp latency differences: A comparison of single-participant and jackknife-based scoring methods. Psychophysiology 45(2):250–274

Knoblich G, Flach R (2001) Predicting the effects of actions: Interactions of perception and action. Psychological science 12(6):467–472

Kovarski K, Latinus M, Charpentier J, Cléry H, Roux S, Houy-Durand E, Saby A, Bonnet-Brilhault F, Batty M, Gomot M (2017) Facial expression related vmmn: Disentangling emotional from neutral change detection. Frontiers in human neuroscience 11:18

Kujala T, Tervaniemi M, Schröger E (2007) The mismatch negativity in cognitive and clinical neuroscience: theoretical and methodological considerations. Biological psychology 74(1):1–19

Laird JD, Lacasse K (2014) Bodily influences on emotional feelings: Accumulating evidence and extensions of william james’s theory of emotion. Emotion Review 6(1):27–34

Lappe C, Lappe M, Pantev C (2016) Differential processing of melodic, rhythmic and simple tone deviations in musicians-an meg study. Neuroimage 124:898–905

Leitman DI, Sehatpour P, Garidis C, Gomez-Ramirez M, Javitt DC (2011) Preliminary evidence of pre-attentive distinctions of frequency-modulated tones that convey affect. Frontiers in human neuroscience 5:96

Maris E, Oostenveld R (2007) Nonparametric statistical testing of eeg-and meg-data. Journal of neuroscience methods 164(1):177–190

Martínez-Montes E, Hernández-Pérez H, Chobert J, Morgado-Rodríguez L, Suárez-Murias C, Valdés-Sosa PA, Besson M (2013) Musical expertise and foreign speech perception. Frontiers in systems neuroscience 7:84

Neuhoff JG (1998) Perceptual bias for rising tones. Nature 395(6698):123

Niedenthal PM (2007) Embodying emotion. science 316(5827):1002–1005

Oostenveld R, Fries P, Maris E, Schoffelen JM (2011) Fieldtrip: Open source software for advanced analysis of meg, eeg, and invasive electrophysiological data. Computational Intelligence and Neuroscience 2011:1

Pakarinen S, Sokka L, Leinikka M, Henelius A, Korpela J, Huotilainen M (2014) Fast determination of mmn and p3a responses to linguistically and emotionally relevant changes in pseudoword stimuli. Neuroscience letters 577:28–33

Pampalk E (2004) A matlab toolbox to compute music similarity from audio. In: ISMIR

Pannese A, Hirsch J (2011) Self-face enhances processing of immediately preceding invisible faces. Neuropsychologia 49(3):564–573

Peirce JW (2007) Psychopy—psychophysics software in python. Journal of neuroscience methods 162(1):8–13

Pinheiro AP, Barros C, Dias M, Niznikiewicz M (2017a) Does emotion change auditory prediction and deviance detection? Biological psychology 127:123–133

Pinheiro AP, Barros C, Vasconcelos M, Obermeier C, Kotz SA (2017b) Is laughter a better vocal change detector than a growl? Cortex 92:233–248

Rachman L, Liuni M, Arias P, Lind A, Johansson P, Hall L, Richardson D, Watanabe K, Dubal S, Aucouturier JJ (2018) David: An open-source platform for real-time transformation of infra-segmental emotional cues in running speech. Behavior Research Methods 50(1):323–343

Rohr L, Abdel Rahman R (2015) Affective responses to emotional words are boosted in communicative situations. NeuroImage 109:273–282

Sato M, Tremblay P, Gracco VL (2009) A mediating role of the premotor cortex in phoneme segmentation. Brain and language 111(1):1–7

Schall S, Kiebel SJ, Maess B, von Kriegstein K (2015) Voice identity recognition: Functional division of the right STS and its behavioral relevance. Journal of Cognitive Neuroscience 27(2):280–291, DOI 10.1162/jocn

Schirmer A, Striano T, Friederici AD (2005) Sex differences in the preattentive processing of vocal emotional expressions. Neuroreport 16(6):635–639

Schirmer A, Simpson E, Escoffier N (2007) Listen up! processing of intensity change differs for vocal and nonvocal sounds. Brain research 1176:103–112

Scott SK, Johnsrude IS (2003) The neuroanatomical and functional organization of speech perception. Trends in neurosciences 26(2):100–107

Sel A, Forster B, Calvo-Merino B (2014) The emotional homunculus: Erp evidence for independent somatosensory responses during facial emotional processing. Journal of Neuroscience 34(9):3263–3267

Sel A, Harding R, Tsakiris M (2016) Electrophysiological correlates of self-specific prediction errors in the human brain. Neuroimage 125:13–24

Skipper JI, Devlin JT, Lametti DR (2017) The hearing ear is always found close to the speaking tongue: Review of the role of the motor system in speech perception. Brain and Language 164:77–105

Sulykos I, Kecskés-Kovács K, Czigler I (2015) Asymmetric effect of automatic deviant detection: the effect of familiarity in visual mismatch negativity. Brain research 1626:108–117

Tacikowski P, Nowicka A (2010) Allocation of attention to self-name and self-face: An ERP study. Biological Psychology 84(2):318–324

Tadel F, Baillet S, Mosher JC, Pantazis D, Leahy RM (2011) Brainstorm: a user-friendly application for meg/eeg analysis. Computational intelligence and neuroscience 2011:8

Tye-Murray N, Spehar BP, Myerson J, Hale S, Sommers MS (2015) The self-advantage in visual speech processing enhances audiovisual speech recognition in noise. Psychonomic bulletin & review 22(4):1048–1053

Ulrich R, Miller J (2001) Using the jackknife-based scoring method for measuring lrp onset effects in factorial designs. Psychophysiology 38(5):816–827

Wagenmakers EJ, Beek T, Dijkhoff L, Gronau QF, Acosta A, Adams Jr R, Albohn D, Allard E, Benning S, Blouin-Hudon EM, et al (2016) Registered replication report: Strack, martin, & stepper (1988). Perspectives on Psychological Science 11(6):917–928

Wilson SM, Iacoboni M (2006) Neural responses to nonnative phonemes varying in producibility: Evidence for the sensorimotor nature of speech perception. Neuroimage 33(1):316–325

